# Design of an effective sgRNA for CRISPR/Cas9 knock-ins in polyploid *Synechocystis sp*. PCC 6803

**DOI:** 10.1101/2023.09.25.559380

**Authors:** María Isabel Nares-Rodriguez, Esther Karunakaran

**Affiliations:** Department of Chemical and Biological Engineering, University of Sheffield, Sheffield, United Kingdom

**Keywords:** CRISPR/Cas9, sgRNA design, mutant segregation, *Synechocystis sp*. PCC 6803, minimal induction

## Abstract

*Synechocystis sp*. PCC 6803 (*Synechocystis*) is a highly promising organism for the production of diverse recombinant chemicals, including biofuels. However, conventional genetic engineering in *Synechocystis* is challenging due to its highly polyploid genome which not only leads to low product yields but also makes the recombinant organism less reliable for use in biomanufacturing. Due to its precision, effectiveness and reliability in a vast array of chassis, CRISPR/Cas9 has the potential of overcoming the drawbacks effected by a polyploid genome. Here we identified and developed an effective sgRNA for the knock-in of nucleotide sequences of varying lengths in the neutral site *slr*0168 of polyploid *Synechocystis* using CRISPR/Cas9. The gene encoding digeranylgeranylglycerophospholipid reductase from *Sulfolobus acidocaldarius* and the methyl ketone operon from *Solanum habrochaites* were chosen as the exemplar nucleotide sequences for incorporation into the chromosome of *Synechocystis*. It is demonstrated here that our sgRNA design was effective for both knock-ins and that CRISPR/Cas9 achieves complete mutant segregation after a single step of selection and induction.

## 1. Introduction

An important aim of genetic engineering is the expression of heterologous proteins in a host organism that enables the generation of a target product. For robust industrial production, the genes encoding these heterologous proteins must be stable and homogenic in the host, that is, the genes must be borne on all chromosomal copies so that they are inherited and maintained in every descendant in each cell-division cycle to prevent mutation dilution and mutant loss during manufacturing and ensure the high yields of the desired product (1-4).

The polyploid model cyanobacterium *Synechocystis sp*. PCC 6803 (hereafter *Synechocystis*) is a highly promising chassis for the production of many recombinant products relevant to a number of industries, including biofuels and personal care products (5-10). However, traditional genetic engineering of *Synechocystis* is laborious, time-consuming and not cost-effective due to its highly polyploid genome. To overcome this drawback, strategies such as phosphate deprivation, multiple counter-selection and combined random mutagenesis have been implemented (11-13). These strategies rely on the natural occurring mechanism of double homologous recombination (DHR) in *Synechocystis* (7, 12, 14, 15) and full mutant segregation remains difficult to achieve (11-13). Consequently, gene expression as well as target product yields are often compromised precluding use of *Synechocystis* as the host in the industrial-scale production of heterologous target products (4, 16-18). However, emerging technologies, such as CRISPR/Cas9 may alleviate this bottleneck.

The type II CRISPR system (clustered regularly interspaced short palindromic repeats) and its associated nuclease Cas9 (CRISPR/Cas9) is an adaptive immune system in Archaea and Bacteria that has been adapted for highly precise and effective genome editing. The CRISPR/Cas9 system implements a single guide RNA (sgRNA), a Cas9 nuclease and a donor DNA (dDNA) to introduce the latter with high precision in a target locus in the host’s genome via its natural homology directed repair mechanism triggered by the double strand breaks (DSB) effected by Cas9 (19-23). Previous gene editing strategies are based on DNA-protein interactions which are considered highly complex. These have not reached the level of effectiveness and precision deemed to CRISPR/Cas9 (23-26), which acts upon nucleotide sequence complementarity and is considered to be simpler; this endows CRISPR/Cas9 with its highly elevated rate of success (20, 27, 28). Moreover, CRISPR/Cas9 is considered to be efficient, reliable and precise due to its chimeric and easily reprogrammable sgRNA, activity of Cas9 and the repair mechanism activated by DSB (29-31). CRISPR/Cas9 is able to perform multiplexed insertions, inactivation, deletions and editing of genes in multiple loci in a simultaneous manner (20, 32-34) and has successfully been implemented in a wide range of hosts, including humans (25, 29, 30, 35-38). Therefore, CRISPR/Cas9 may have the potential of overcoming bottlenecks such as the polyploid genome in *Synechocystis*.

Thus, with the hypothesis that CRISPR/Cas9 can favour mutant segregation, we engineered polyploid *Synechocystis* via CRISPR/Cas9 by separately knocking-in heterologous digeranylgeranylglycerophospholipid reductase (GGR) and a methyl ketone operon (ShMKS), to evaluate 1) if CRISPR/Cas9 can indeed improve mutant segregation, 2) if the size and complexity of the dDNA (GGR-dDNA (single gene) or ShMKS-dDNA (dual-gene operon)) has an impact on CRISPR/Cas9 and mutant segregation and 3) if the CRISPR/Cas9 effects would vary if applied upon markerless and selection marker-bearing dDNA. We implemented an empirical method based on available tools and principles to identify and design an effective sgRNA for successfully inserting the desired genes in the *slr*0168 neutral site, achieving full mutant segregation after a single step of induction and selection for both, markerless as well as mutants bearing a selection marker in the dDNA.

## 2. Materials and methods

### 2.1 Strains and media

*E. coli* 10-beta was used as a cloning strain. The cells were cultivated at 37 °C in sterile lysogeny broth (LB) medium at an orbital agitation of 150 rpm for aeration. The strain was used for all the molecular cloning experiments, plasmid propagation and maintenance. *E. coli* copy cutter cells bearing the pPMQK1 vector were cultivated at 37 °C in sterile LB medium at an orbital agitation of 150 rpm for aeration. The medium was supplemented with kanamycin, at a concentration of 50 μg/mL. The strain was used for the propagation of the plasmid bearing the Cas9 gene. *E. coli* MC1061 cells were cultivated at 37 °C in sterile LB medium at an orbital agitation of 150 rpm for aeration. The strain was used as a helper strain in all tri-parental mating events. The medium was supplemented with chloramphenicol at a concentration of 25 μg/mL and ampicillin at a concentration of 100 μg/mL for plasmid maintenance where appropriate.

The *Synechocystis* wild type (WT) strain (*Synechocystis sp*. PCC 6803) used for the CRISPR mutagenesis was purchased from the Pasteur Culture Collection of Cyanobacteria (PCC). All *Synechocystis* cells were grown photoautotrophically in sterile Blue-Green liquid medium (BG-11) at a temperature of 25°C, under continuous orbital agitation conditions of 100 rpm for aeration. Initially, the mutants were placed under white light, with an intensity of 125 μmol/m2/s; subsequently, this was changed to red light, with an intensity of 20 μmol/m2/s. The media for *Synechocystis* mutants bearing chloramphenicol resistance was supplemented with chloramphenicol at a concentration of 12 μg/mL.

### 2.2 Insert and vector construction

The pPMQK1 vector, which was a kind gift from Dr. Paul Hudson and his group (Karolinska Institute; Solna, Sweden) carried the *Streptococcus pyogenes* Cas9 nuclease expression cassette and was transformed into *Synechocystis* for its expression. The pEERM4 Cm (Addgene plasmid # 64026) and the pEERM3 Km vectors (Addgene plasmid # 64025) were kind gifts from Pia Lindberg (39). The QIAGEN Miniprep Spin Kit was used following manufacturer’s instructions for plasmid extraction from *E. coli* strains. For plasmid manipulation, appropriate enzymes were sourced from New England Biolabs (UK) and manufacturer’s instructions were followed. Chemically competent *E. coli* cells were purchased from New England Biolabs (UK) and the heat shock method was method was performed following manufacturer’s instructions for plasmid verification and maintenance. Once designed, the sgRNA and donor DNA sequences were synthesized as custom genes by Eurofins Genomics.

### 2.3 sgRNA selection

All available protospacer candidates in *slr*0168 were identified using the CasOT software (40). To enable the identification of suitable protospacer sequences, the sequence of *slr*0168 was submitted to CasOT as a FASTA file and the following parameters were set as the selection criteria. The -NGG Protospacer Adjacent Motif (PAM) only option (option “A-NGG only”) was selected to generate only candidates to match this PAM. The “target and off-target” mode was selected to allow the generation of all the possible protospacer candidates followed by off-target analysis. A maximum of 2 mismatches were allowed in the seed region. The “non-5’G” requirement option was selected as the T7 promoter was not used in the sgRNA. And, finally, a “length” of 19 to 22 bp for the base-pairing region of the candidates was allowed. For the off-target analysis, each protospacer candidate was evaluated for the number of possible mismatches and off-targets they presented.

### 2.4 Cas9 handle secondary structure analysis

The RNAfold online tool from the ViennaRNA Package was used to evaluate if the Cas9 handle formed a proper secondary structure after the addition of the selected protospacer (41). The selected 20 nt protospacer was appended to the 42 nt sequence of the Cas9 handle, for a total of 62 nt sequence and subsequently input into the algorithm to evaluate its secondary structure formation. The minimum free energy (MFE) and partition function, dangling energies on both sides of a helix, the Turner model (2004) options for RNA parameters and the default criteria for the output options were selected.

### 2.5 Strain construction and CRISPR/Cas9 induction

The pPMQK1 vector was transformed into *Synechocystis* to generate *Synechocystis::*Cas9 through standard protocol for tri-parental mating in cyanobacteria (42). Subsequently, the pEERM4::sgRNA_GGR-dDNA and the pEERM3::ShMKS-dDNA vectors were transformed independently into *Synechocystis::*Cas9 following a standard protocol for tri-parental mating in cyanobacteria (42) to generate *Synechocystis:*:Cas9*_*sgRNA_GGR-dDNA and *Synechocystis*::Cas9_sgRNA_ShMKS-dDNA respectively.

The transformants were transferred to non-selective BG-11 plates for 24 hours for recovery and then subjected to induction and selection. Only ShMKS mutants were subjected to both, selection and induction while GGR mutants were only subjected to induction, effectuated on BG-11 plates supplemented with anhydrotetracycline (aTc) at a concentration of 200 ng/mL while using white light for the first attempt and a subsequent one with an aTc concentration of 400 ng/mL under red light.

### 2.6 Mutant segregation analysis

The segregation analysis was performed on genomic DNA extracts of *Synechocystis* and *Synechocystis* mutants obtained using the Gen-Elute Bacterial Genomic DNA Kit (Sigma-Aldrich) by effectuating a screen for GGR and ShMKS incorporation into *slr*0168 using PCR. The PCR analysis was performed using the GoTaq Green Master Mix (Promega). Segregation primers (Supplementary Table S1) amplifying GGR-dDNA and ShMKS-dDNA from the homology regions were designed and used for analysis. The mutants were screened after a single round of selection/induction to determine their state of mutant segregation.

## 3. Results

### 3.1 Selection of the target site, Cas9 nuclease and protospacer

The gene *slr*0168 was selected as the target site for the incorporation of genes encoding heterologous proteins (cargo). It was chosen as it has been identified as a highly effective neutral site in *Synechocystis* (43) and therefore incorporation of the cargo at this site will avoid any undesired phenotypical changes to the host.

The Cas9 nuclease from *S. pyogenes* was chosen as it is the most widely employed nuclease for genome editing using CRISPR (44, 45) and recognizes an NGG PAM (46, 47), which has been reported to hold the highest activity when compared to other PAMs (48-50). However, the specificity of CRISPR elements, such as the sgRNA towards the target site is not ensured; hence, the protospacer region of the sgRNA could anchor to other sites instead of its target, leading to unintended editing by off-target effects (51, 52).

The protospacer is the sgRNA region that is complementary to the target site within the genome. Hence, this section is designed to anchor to the region adjacent to the PAM in order to allow the cleavage in the target to be edited (53). Within a protospacer, the first 12 nt closer to the PAM encompass the seed region, critical for proper cleavage. Mismatches in such region could significantly reduce the cleavage activity of the sgRNA/Cas9 complex while mismatches in the non-seed region (rest of protospacer) might have a less critical impact but can still increase the probabilities of off-target effects (48, 49). Hence, the fewer number of mismatches within the seed region would be less likely to have an impact on cleavage activity. Similarly, the fewer number of both, overall missmatches and off-targets would increase the probabilities of effective target-binding from a protospacer; which is why careful evaluation is crucial in order to identify the probable off-target activity of a given protospacer to increase its specificity and cleavage activity.

Therefore, using the CasOT software (40), the target gene was scanned in order to identify all the available protospacer candidates associated with the designated PAM and evaluate their off-target effects. For each candidate, a number of specific types of mismatches represented by codes (e.g. A205, A206) was given, followed by the number of off-targets for each type of mismatch (shaded in blue) (Figure 1).The letter “A” on each code represents the selection of an NGG PAM. The first number after this letter indicates the number of mismatches in the seed region, the following two numbers represent the number of mismatches in the non-seed region (40). A total of 73 protospacer candidates were identified within *slr*0168. The selected protospacer (Supplementary Table S2) which was 20 nt long and within the suggested length for effective binding (22, 53), showed a total of 3 off-target sequences, one off-target generated by the A205 mismatch type and 2 off-targets generated by the A206 mismatch type (Figure 1). Hence, the one identified off-target sequence is shown for the A205 type and 2 sequences are shown for the A206 type (Figure 1). The A205 type indicates two mismatches in the seed region and five in the non-seed region while the A206 type represents 2 mismatches on the seed region and 6 in the non-seed region. The A000 code represents the original protospacer candidate, hence, it is not considered neither a type of mismatch nor an off-target. All identified off-target sequences for the selected protospacer are shown (Figure 1).

**Figure 1.**
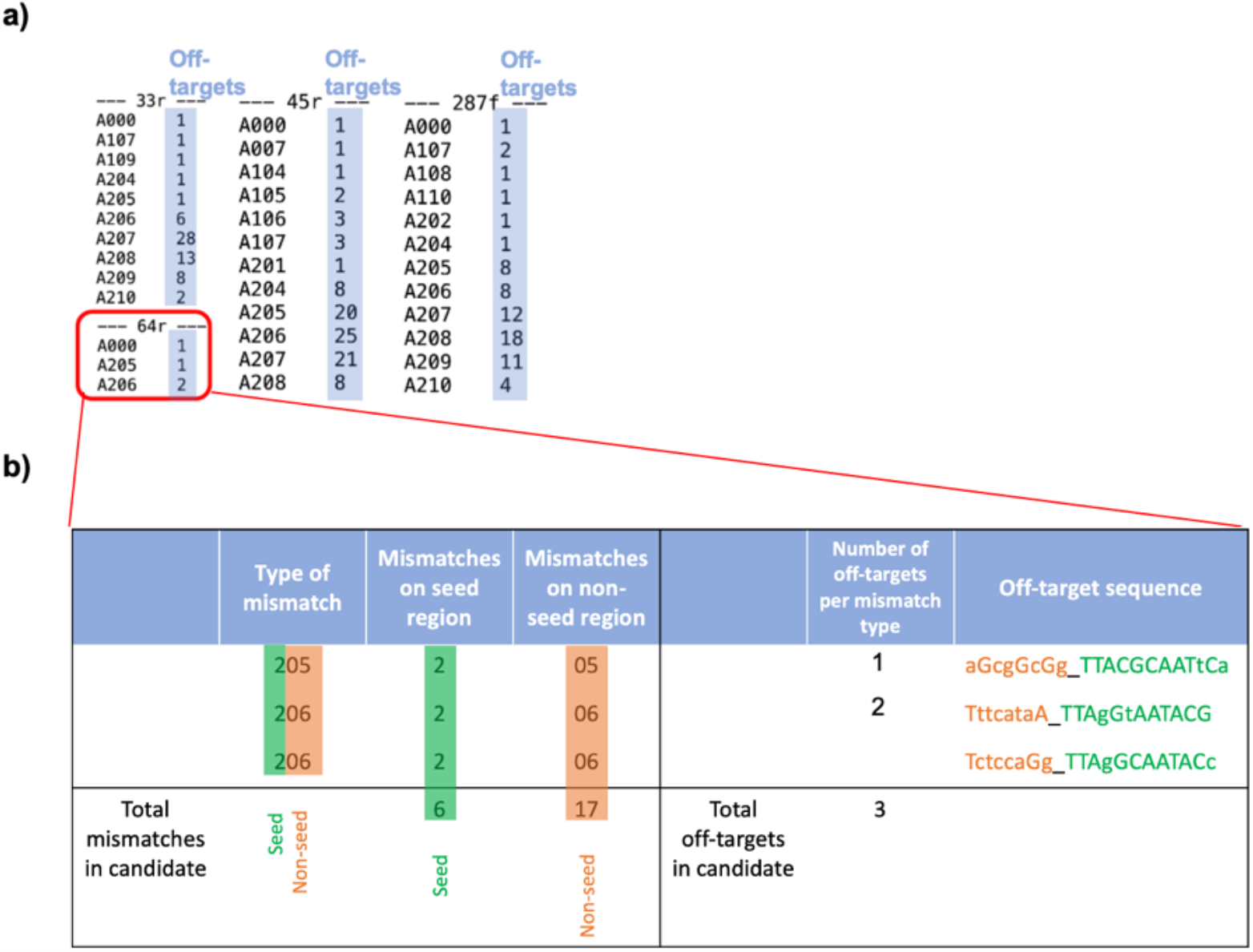
Off-target analysis of the non-selected and selected (in red) protospacers showing the type and number of mismatches (codes; i.e. A206) and number of off-targets for each (shaded in blue) type of mismatch (a). Description of the off-targets and mismatches (lowercase) of the selected protospacer within the seed (green) and the non-seed (orange) region (b).

Compared to other candidates, the selected protospacer showed the fewest number of both off-targets as well as mismatches. The selected protospacer showed only two types of mismatches (A205 and A206) as opposed to other candidates such as 287f in which up to 11 types of mistatches were identified (A204 to A210). Similarly, only 3 off-targets were identified within the selected protospacer (1 for A205 and 2 for A206) whereas in other candidates such as 33r up to 28 off-targets were identified for a single type of mismatch (A207). The selected protospacer showed the lowest number of mismatches in both, the seed and non-seed region as well as the smallest number of off-targets. Therefore, it was considered the least probable to anchor to off-target sites, highly expected to present the highest activity and enable the knock-ins in the desired target editing site.

The selected protospacer, after integration with the Cas9 handle, must enable the formation of the proper secondary structure on the handle, which subsequently will aid in creating the sgRNA-Cas9 complex to effect the DSB at the target site (20, 41). The sequence of the protospacer contributes to whether the proper secondary structure will be formed within the handle. Moreover, the free energy between the interaction of such elements has, as well, been reported to have an impact in the formation of the proper hairpin. Protospacer-handle sequences showing a free energy value within 5% of the optimal folded structure have been deemed acceptable (41). Therefore, the secondary structure of the selected protospacer was then evaluated to determine if it would enable the formation of a proper handle loop. RNA secondary structure analysis suggested that the selected protospacer enabled the proper hairpin within the handle; moreover, it showed a free energy value of -14.61 kcal/mol and within 5% of the optimal folded structure (-14.30 kcal/mol) (Figure 2). Hence, an effective sgRNA-Cas9 complex is estimated to be formed intracellularly. Conjoined with the other selection factors previously mentioned, the designed sgRNA (Figure 3d) (full sequences are provided in Supplementary Table S2) presented the highest probability of effectiveness for genome editing at *slr*0168.

**Figure 2.**
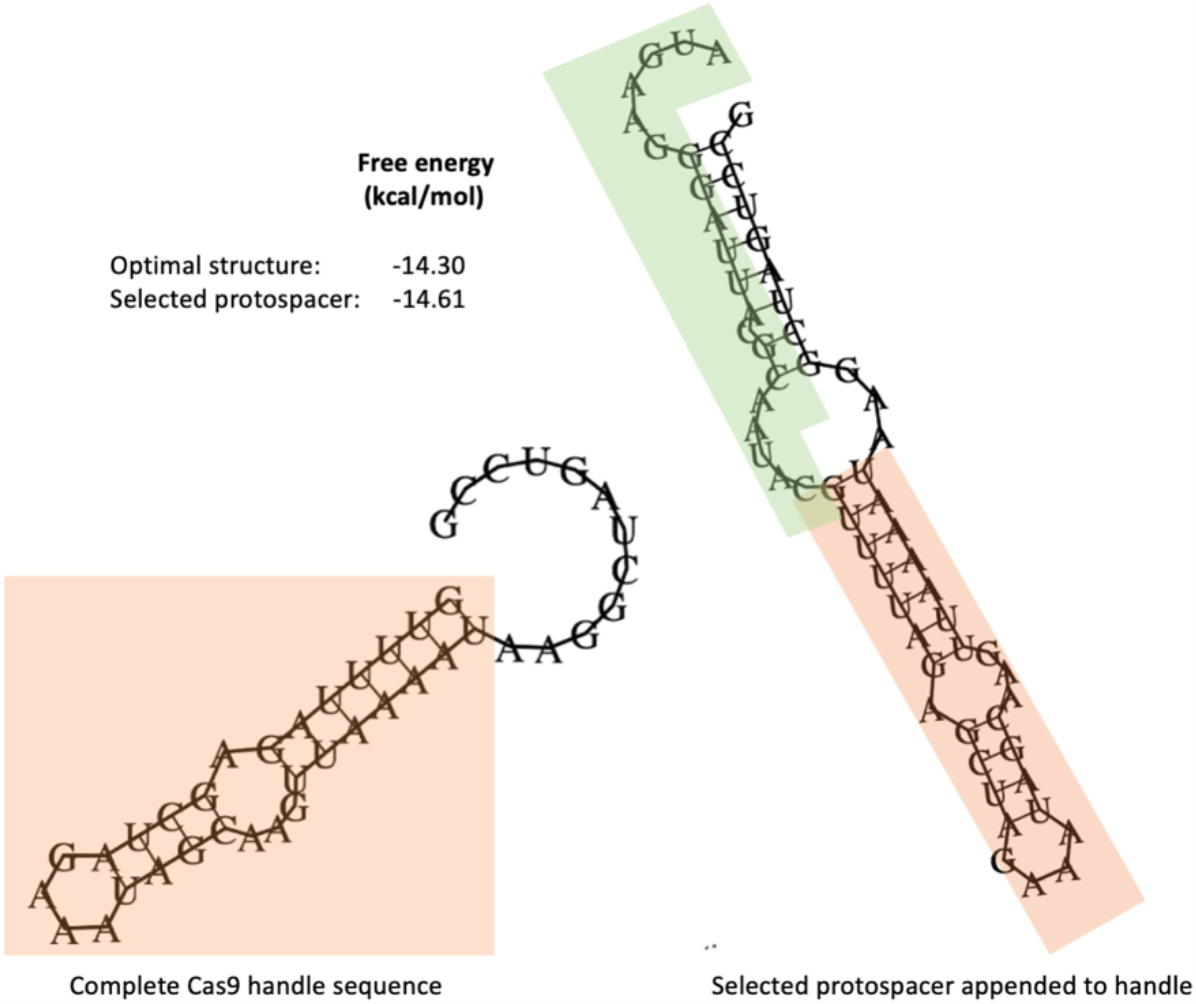
Free energy values and Cas9-handle secondary structure analysis showing the proper structure (shaded in orange) within the complete Cas9 handle sequence and the proper structure also present (shaded in orange) with the selected protospacer appended (shaded in green) to the handle.

**Figure 3.**
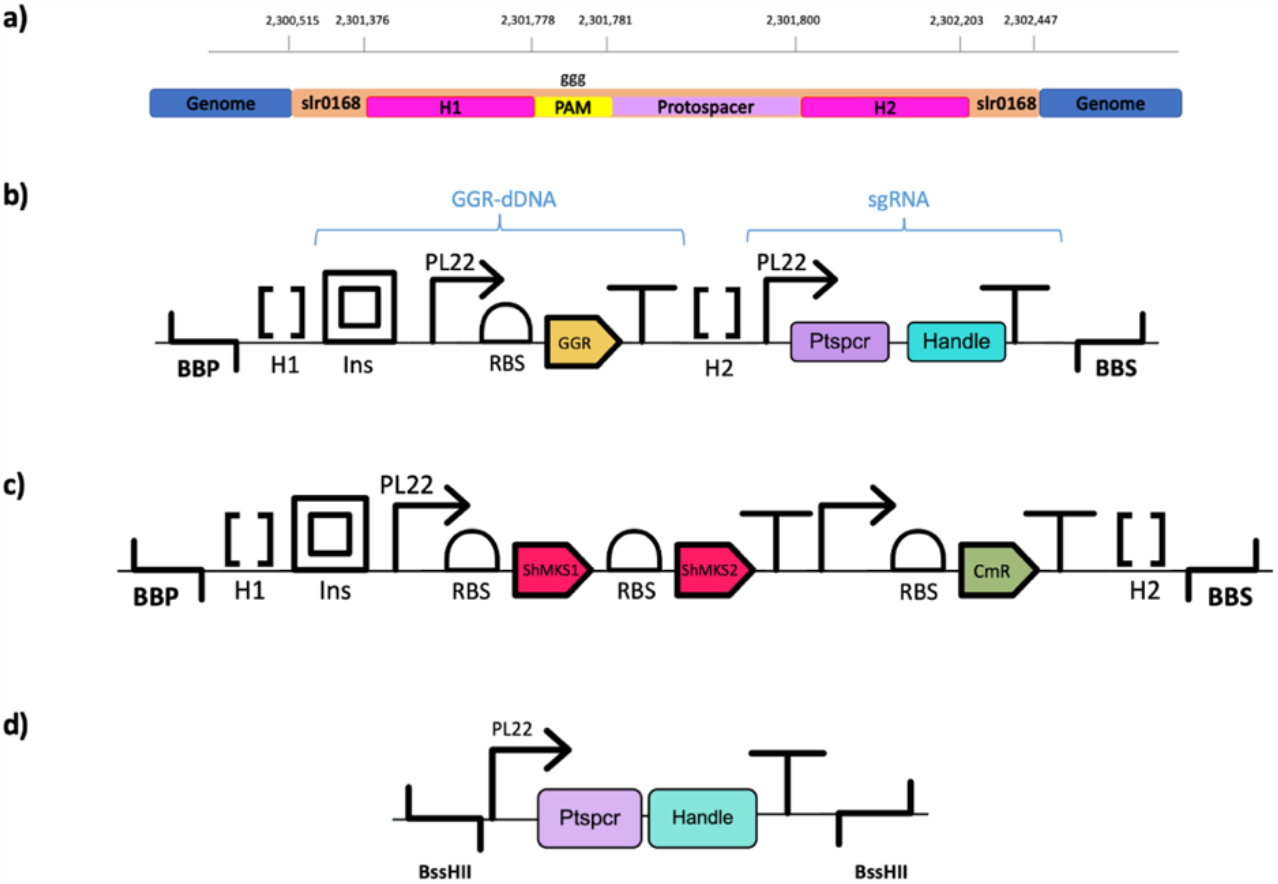
Genetic context of the selected protospacer of the sgRNA, showing elements upstream and downstream of the protospacer in the genome of *Synechocystis* WT (a). Schematic of the sgRNA_GGR-dDNA, including flanking BioBrick prefix and suffix (BBP and BBS), upstream and downstream homology regions (H1 and H2), an insulator (ins), PL22 promoter, selected RBS, heterologous GGR gene (GGR), T7 terminator followed by the sgRNA (b). Schematic illustrating the ShMKS-dDNA similar elements to (b) but two heterologous genes (ShMKS1 and ShMKS2) encompassing the operon and a chloramphenicol resistance marker (CmR) (c). Schematic showing the designed sgRNA, with BssHII overhangs, PL22 promoter, selected protospacer (Ptspcr), Cas9 handle and *S. pyogenes* terminator (d).

### 3.2 Design of the donor DNA templates

Two donor DNA templates were designed in this study, the sgRNA_GGR-dDNA (Figure 3b). and the ShMKS-dDNA (Figure 3c). One of the templates encompassed the coding sequence corresponding to the enzyme digeranylgeranylglycerophospholipid reductase (GGR) from *Sulfolobus acidocaldarius* (Genbank accession # SACI_RS04710; 1,359 bp) whilst the other template encompassed the coding sequences corresponding to enzymes MKS1 and MKS2 belonging to the methyl ketone synthesis operon from *Solanum habrochaites* (Genbank accession/ENA coding Id # AAV87156.1; 798 bp and # ADK38536.1; 627 bp, respectively). The coding sequences in both templates were placed under the control of the anhydrotetracycline-inducible PL22 promoter, the B0034 RBS and the T7 terminator. PL22 was chosen as it has been reported to provide a suitable expression of CRISPRi elements (54, 55). In both templates, the assembled genetic circuits were placed in between homology arms comprising approximately 400 bp for the GGR mutants and approximately 200 bp for the ShMKS mutants, upstream and downstream of the chosen protospacer sequence in *slr*0168 (Figure 3a, exact base positions on the genome are provided in Supplementary Table S3 and S4). Care was taken to omit the PAM sequence from inclusion in the homology arms to prevent unintended and continuous cleavage from Cas9 at the target site.

Once the basic design of the donor DNA was complete, two divergent approaches were taken with respect to the further design. The template harbouring the GGR was chosen arbitarily to generate a markerless knock-in mutant for the evaluation of Cas9-based genome editing in *Synechocystis*. Therefore an antibiotic selection marker was not included in the template. The total size of the cargo including the homology arms was 2,293 bp and was titled sgRNA_GGR-dDNA. The genetic circuit harbouring the coding sequence for the designed sgRNA under the control of the PL22 promoter and the *S. pyogenes* terminator was placed in the vicinity of the cargo. The assembly was placed in the pEERM4 plasmid backbone. Conversely, the template harbouring the MKS1 and MKS2 coding sequences was chosen to generate a knock-in mutant with chloramphenicol resistance as the selection marker. The assembly was placed in the pEERM3 plasmid backbone. The total size of the cargo including the homology arms was 2,936 bp and was titled ShMKS-dDNA. The genetic circuit harbouring the coding sequence for the designed sgRNA under the control of the PL22 promoter in this template was inserted into the BssHII site of the backbone.

*Synechocystis* bearing the gene encoding the *S. pyogenes* Cas9 on plasmid pPMQK1 under the control of the PL22 promoter was used as the host organism. This strain was independently transformed with either donor DNA template and subjected to a single round of induction with anhydrotetracycline.

### 3.3 Mutant segregation analysis

Induction of the CRISPR/Cas9 system was carried out in both strains with an aTc concentration of 200 ng/mL and white light for a first attempt and subsequently modified to 400 ng/mL of aTc and red light for a second attempt. The segregation state of the mutants was evaluated by PCR screening after a single round of selection/induction. For the first attempt, neither mutants showed full mutant segregation (data not shown). However, full mutant segregation was achieved on the second attempt after a single round of selection/induction for *Synechocystis:*:Cas9*_*sgRNA_GGR-dDNA (Figure 4a) as well as for *Synechocystis:*:Cas9*_*sgRNA_ShMKS-dDNA (Figure 4b). The mutant segregation analysis shows a band of approximately 350 base pairs for *Synechocystis* wild type strain (WT). This band corresponds to the homology regions (H1 and H2) flanking GGR-dDNA as well as ShMKS-dDNA and taken from the target neutral site *slr*0168 in the genome of *Synechocystis* wildtype. Therefore, this band, is expected to appear in the wildtype genome. Moreover, the amplicon from the pEERM4 vector bearing the GGR-dDNA shows a band of approximately 1,800 bp, corresponding to the sequence of the GGR-dDNA. Similarly, the amplicon from the pEERM3 vector bearing the ShMKS-dDNA shows a band of approximately 2,800 bp, corresponding to the sequence of the ShMKS-dDNA. Both bands are not only expected since they are positive controls but also, represent the desired bands aimed to be present in the analysis of both mutants, respectively.

**Figure 4.**
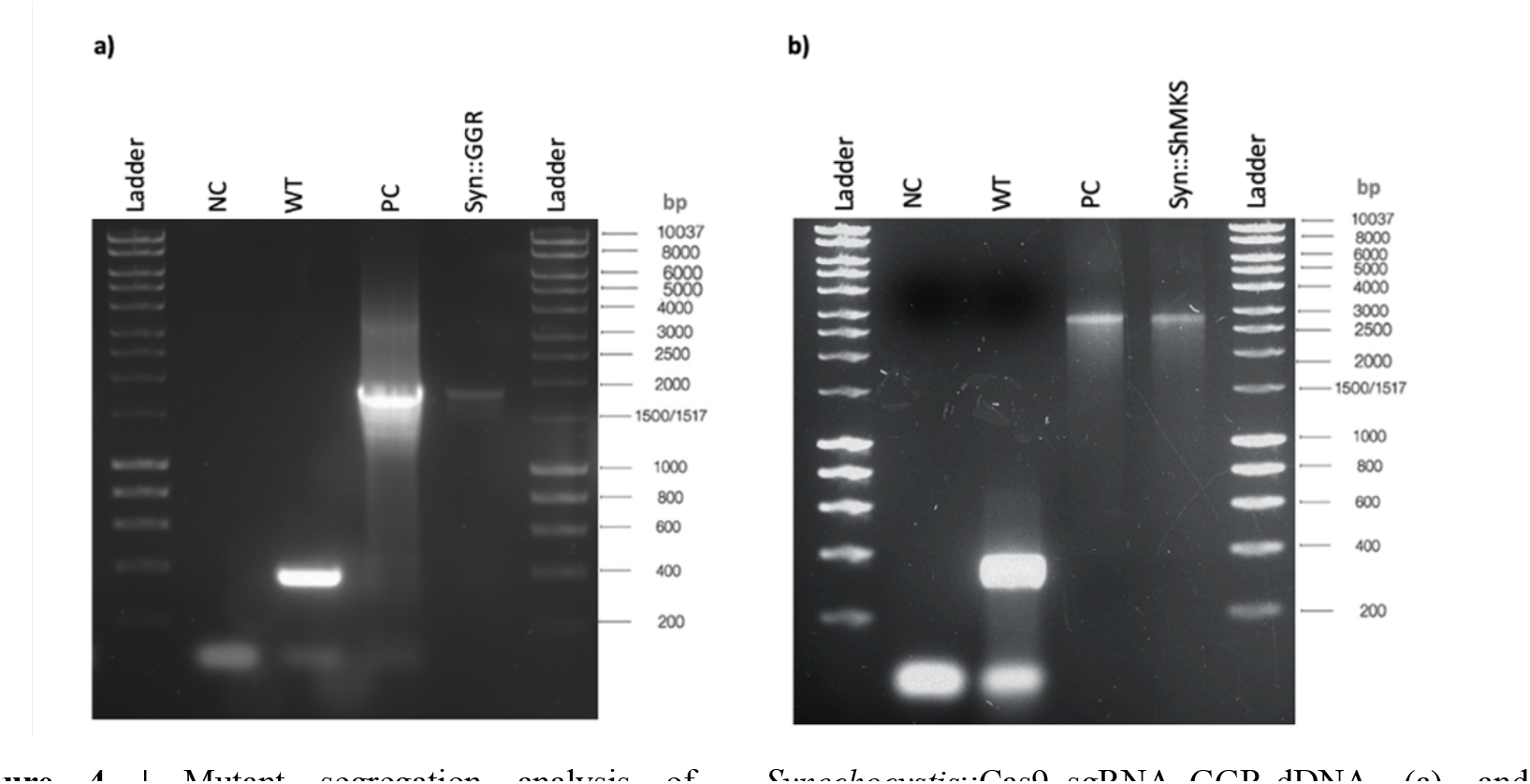
Mutant segregation analysis of *Synechocystis*::Cas9_sgRNA_GGR-dDNA (a) and *Synechocystis::*Cas9_sgRNA_ShMKS-dDNA (b). Showing a negative control (NC); *Synechocystis* wild type (WT) displaying a band corresponding to the homology regions present in the GGR-dDNA, and ShMKS-dDNA taken from the target site in the genome of *Synechocystis* WT; a positive control (PC), displaying a amplicon of the desired size; *Synechocystis*::Cas9_sgRNA_GGR-dDNA for (a) and *Synechocystis*::Cas9_sgRNA_shMKS-dDNA for (b), both showing a band of the desired size that coincides with the desired band on their respective positive control as well as the lack of additional bands for both, (a) and (b).

Furthermore, the mutant segregation analysis for *Synechocystis*::Cas9_sgRNA_GGR-dDNA (Figure 4a) showed a band of the desired size which coincides with the desired band on the positive control. This suggests that the GGR-dDNA sequence is present in the genome of such mutants and, hence, that knock-in of the GGR-dDNA via CRISPR/Cas9 was effective. Furthermore, the desired band is present as a single band, highlighting the absence of additional bands. This indicates that only complete sequences of the GGR-dDNA were found in this mutant population and that the insertion of the GGR-dDNA into all chromosomal copies of all the individuals of the *Synechocystis*::Cas9_sgRNA_GGR-dDNA population was successful. It must be highlighted that this screening was made after a single round of induction. According to these results, the design and selected sgRNA was effective in the knock-in of a single gene up to 1,455 bp. Placing the sgRNA in close vicinity to the cargo did not affect the incorporation of the cargo at *slr*0168. Furthermore, mutant segregation does not seem to be affected although the template DNA did not bear a selection marker.

Comparably, the mutant segregation analysis for *Synechocystis*::Cas9_sgRNA_ShMKS-dDNA (Figure 4b), shows, as well, that the expected band of the desired size is present for the CRISPR/Cas9 mutant which coincides with the desired band on the positive control. Furthermore, the desired band is present as a single band in the lane and there are no other additional bands, similar to the GGR mutant. The results indicate the insertion of the ShMKS-dDNA into all chromosomal copies of *Synechocystis* was successful. Thus, only complete sequences of the ShMKS-dDNA are present in all chromosomal copies within each cell and that all of these mutants in the culture bear the insert in their genome. It must be highlighted that this screening was made after a single round of antibiotic selectection/induction and that the ShMKS mutant bears a dual operon construct and a resistance marker gene, considerably larger than the GGR mutant. According to these results, the size and complexity of the dDNA, up to 2,399 bp, appears to have no effects on CRISPR/Cas9’s ability of favouring mutant segregation after a single round of selection/induction. Furthermore, mutant segregation does not seem to be affected when the sgRNA and dDNA are constructed as distinct inserts in the same vector for inserts bearing a selection marker.

## 4. Discussion

Genetic engineering aims to express heterologous desired traits to ultimately generate industrial scale yields of a target product in a pre-determined chassis. To achieve this, the mutant culture must be fully segregated. The introduced traits must be homogenic and stable, they must be present in all chromosome copies of an individual and in all individuals. Failing to ensure the presence of the desired mutation in all chromosomal copies of all individuals will lead to mutation dilution, ultimately leading to loss of the mutants and low to null yields of the desired product (1, 3). Therefore, full mutation segregation is pivotal for industrial scale biomanufacturing. Furthermore, the achievement of full mutant segregation is more complicated in polyploid genomes such as *Synechocystis* (3, 56, 57). Nevertheless, CRISPR/Cas9 has the potential of overcoming this drawback given it has been deemed highly effective and successful (27, 30).

In this study, an efficacious sgRNA was designed for the successful knock-in of heterologous genes up to 2,399 bp long in the neutral site *slr*0168 of the highly polyploidy *Synechocystis* chassis with complete mutant segregation after a single round of selection/induction through CRISPR/Cas9. The selected protospacer showed the least number of both, mismatches as well as off-targets, resulting in effective insertions at the target site by the designed sgRNA. Similarly, the selection of a sgRNA with the proper secondary structure formation of the Cas9handle constitutes another important factor to be considered in order to allow the correct formation of the sgRNA-Cas9 complex intracellularly and enable the DSB.

In the same fashion, it is important to highlight the use of red light, as opposed to white, when selecting the PL22 promoter for driving expression. The expression of all of the CRISPR/Cas9 elements was controlled by the aTc-inducible PL22. In the first attempt, white light was implemented and full mutant segregation was not achieved. This might have probably been caused by the sensitivity of aTc to white light (58). And even though PL22 has been deemed to drive reasonable expression with CRISPRi, it has as well described as weak under white light in some instances (59). Therefore, it is evidenced here that red light drives the best expression of PL22 in *Synechocystis*. Moreover, we designed all of the CRISPR/Cas9 elements to be driven by the same promoter in order to tune their induction under a single inducer and minimize interference. On the same token, we have as well demonstrated that the increased concentration of aTc to 400 ng/mL is more effective than the original of 200 ng/mL, which is why we recommend such concentration when working with PL22, red light, CRISPR/Cas9 and *Synechocystis*.

For both knock-ins, our designed sgRNA was able to achieve full mutant segregation after a single round of selection/induction in the mutagenesis mediated by CRISPR/Cas9. According to these results, the size and complexity of the dDNA appears to have no effects on the ability of the constructed CRISPR/Cas9 systems to achieve complete mutant segregation. Operons up to 2 genes and a resistance marker gene, with an insert size close to 2,399 bp appears to be easily managed by the CRISPR/Cas9 machinery regarding full mutant segregation after a single round of selection/induction. Furthermore, mutant segregation is highly favoured and is achieved to a full state after a single round of selection/induction whether the CRISPR/Cas9 elements are in close proximity to the cargo on the same plasmid or not. Moreover, the efficiency of our designed sgRNA is not affected by the generation of either markerless or mutants bearing a selection marker as it was effective in both instances. Furthermore, the selected length of homology regions of 400 bp and 200 bp seem to have facilitated well the recombination events (60, 61) and favoured CRISPR/Cas9. Similarly, triparental mating technique (42) for all CRISPR/Cas9 transformation events with *Synechocystis* facilitates well the delivery of all the genetic elements.

In conclusion, this work has described the development of an effective sgRNA, able to effectuate knock-ins of heterologous genes up to 2,399 bp in the neutral site *slr0168* of *Synechocystis* driven by CRISPR/Cas9 and achieve full mutant segregation after a single step of selection/induction. This sgRNA will be an effective addition to the Synthetic Biology toolkit for the genome editing of the highly promising but highly polyploid host organism, *Synechocystis*.

