## Supplementary Tables for "Design of an effective sgRNA for CRISPR/Cas9 knock-ins in polyploid *Synechocystis sp*. PCC 6803"

| *Supplementary Table S1 – Primers used for segregation analysis* | |
| --- | --- |
| *Segregation primers:* | ***Forward (5’ to 3’):***  *atcctcaaaggggacgaagccg*  ***Reverse (5’ to 3’):***  *tggcttggtcatcagcggc* |

| ***Supplementary Table S2*** *- Sequences of the elements of the sgRNA* | |
| --- | --- |
| **Element** | **Nucleic acid sequence(5’ to 3’)** |
| ***PL22 promoter*** | *tccctatcagtgatagagattgacatccctatcagtgatagagatactgggagcta* |
| ***Protospacer*** | *atgaagggattacgcaatac* |
| ***Cas9 handle*** | *gttttagagctagaaatagcaagttaaaataaggctagtccg* |
| ***Terminator from S. pyogenes*** | *ttatcaacttgaaaaagtggcaccgagtcggtgcttttttt* |

| ***Supplementary Table S3*** *- Accession numbers, bp position or sequences of the elements of the*  *sgRNA_GGR-dDNA* | |
| --- | --- |
| Element | **Accession ID, position in the genome or nucleic acid sequence (5’ to 3’)** |
| *H1* | *Start: 2,301,376. End: 2,301,776 (bp end and start position in the genome of Synechocystis)* |
| *Insulator* | *tctagcatgttaaactaga* |
| *PL22 promoter* | *tccctatcagtgatagagattgacatccctatcagtgatagagatactgggagcta* |
| *Ribosome binding site(RBS)* | *BBa_B0034 (iGeM ID)* |
| *GGR* | *SACI_RS04710 (Genbank accession number)(search as “Gene”)* |
| *T7*  *Terminator* | *ggctcaccttcgggtgggcctttctgcg* |
| *H2* | *Start: 2,301,803. End: 2,302,203 (bp end and start position in the genome of Synechocystis)* |
| *PL22 promoter* | *tccctatcagtgatagagattgacatccctatcagtgatagagatactgggagcta* |
| *Protospacer* | *tgaagggattacgcaatac* |
| *Cas9 handle* | *gttttagagctagaaatagcaagttaaaataaggctagtccg* |
| *Terminator from S. pyogenes* | *ttatcaacttgaaaaagtggcaccgagtcggtgctttttt* |
| *Slr0168 gene* | *Start: 2,300,515. End: 2,302,447 (bp end and start position in the genome of Synechocystis)* |
| *BioBrick prefix* | *gaattcgcggccgcttctagag* |
| *BioBrick suffix* | *tactagtagcggccgctgcag* |

| *Supplementary Table S4 - Accession numbers, bp position or sequences of the elements of the ShMKS-dDNA* | |
| --- | --- |
| Element | **Accession ID, bp position or sequence (5’ to 3’)** |
| *BioBrick prefix (BBp)* | *gaattcgcggccgcttctagag* |
| *H1* | *Start: 2,301,542. End: 2,301,776 (bp end and start position in the genome of Synechocystis)* |
| *Insulator* | *agctgtcaccggatgtgctttccggtctgatgagtccgtgaggacgaaacagcctctacaaataattttgtttaaactaga* |
| *PL22 promoter* | *tccctatcagtgatagagattgacatccctatcagtgatagagatactgggagcta* |
| *Ribosome binding site (RBS)* | *BBa_B0034 (iGeM ID)* |
| *ShMKS1* | *AAV87156.1 (ENA coding Id)* |
| *ShMKS2* | *ADK38536.1 (ENA coding Id)* |
| *T7*  *Terminator* | *ggctcaccttcgggtgggcctttctgcg* |
| *CmR* | *Chloramphenicol resistance cassette from plasmid pEERM4 (Addgene plasmid # 64026)* |
| *H2* | *Start: 2,301,803. End: 2,302,015 (bp end and start position in the genome of Synechocystis)* |
| *BioBrick suffix (BBs)* | *tactagtagcggccgctgcag* |
